# “The flavor enhancer maltol increases pigment aggregation in dermal and neural melanophores in *Xenopus laevis* tadpoles”

**DOI:** 10.1101/645606

**Authors:** Lara I. Dahora, Ashley Fitzgerald, Matthew Emanuel, Alexa F. Baiges, Zahabiya Husain, Christopher K. Thompson

**Affiliations:** Department of Biological Sciences, Virginia Tech, Blacksburg, VA; School of Neuroscience, Virginia Tech, Blacksburg, VA; Global Change Center, Virginia Tech, Blacksburg, VA

**Keywords:** melanophore, maltol, pigment, aggregation, melatonin

## Abstract

Melanophores are pigmented cells that change the distribution of pigmented melanosomes, enabling animals to appear lighter or darker for camouflage, thermoregulation, and UV-protection. A complex series of hormonal and neural mechanisms regulates melanophore pigment distribution, making these cells a valuable tool to screen toxicants as a dynamic cell type that responds rapidly to the environment. We found that maltol, a naturally occurring flavor enhancer and fragrance agent, induces melanophore pigment aggregation in a dose-dependent manner in *Xenopus laevis* tadpoles. To determine if maltol affects camouflage adaptation, we placed tadpoles into maltol baths situated over either white or black background. Maltol induced pigment aggregation in a similar dose-dependent pattern regardless of background color. We also tested how maltol treatment compares to melatonin treatment and found that the degree of pigment aggregation induced by maltol is similar to treatment with melatonin, but the time course differs significantly. Last, maltol had no effect on mRNA expression of pro-opiomelanocortin or melanin concentrating hormone receptor in the brain, both of which regulate camouflage-related pigment aggregation. Our results suggest that maltol does not exert its effects via the camouflage adaptation mechanism nor via melatonin-based mechanisms. These results are the first to identify a specific toxicological effect of maltol exposure and rules out several mechanisms by which maltol may exert its effects on pigment aggregation.

## INTRODUCTION

Melanophores are pigmented cells used by a wide range of animals for camouflage, UV protection, and thermoregulation. These cells alter their appearance by adjusting the location of melanin pigment within the cell. When the animal needs to appear darker in color, melanin is dispersed throughout the cell. When lighter color is needed, the melanin is transported to the center of the cell, which is known as pigment aggregation. Melanosomes are organelles contained in melanophores that are responsible for synthesizing and transporting melanin within melanophores. Melanophores have been described as smooth muscle cells in disguise, because myosin V is responsible for melanosome transport along actin filaments, with the help of microtubule associated proteins (MAPs) dynein and kinesin (Salim and Ali 2012). Kinesin II and myosin V work together to mediate dispersion, whereas dynein mediates aggregation by facilitating retrograde transport of melanosomes along microtubules and actin filaments (Reilein et al. 1998; Gross et al. 2002).

The African clawed frog, *Xenopus laevis*, has been a key animal model for better understanding melanophore form and function. Changes in pigment distribution in melanophores are mediated by multiple mechanisms. First, melanophores are themselves photosensitive; they express melanopsin-expressing photoreceptor that controls pigment distribution (Provencio et al. 1998). It has been demonstrated previously that melanophores in *Xenopus* tadpoles exhibit a light sensitive response, with melanophore aggregation increasing after exposure to light, and decreasing during darkness (Moriya et al. 1996). Second, photoperiod contributes to changes in melanophore pigment distribution, which largely regulated by changes in circulating levels of melatonin. Initial experiments investigating the role of melatonin in amphibian pigmentation found that changes in melanophore appearance result from changes in environmental illumination, which regulates release of melatonin (Binkley et al. 1988). Onset of night induces release of melatonin from the pineal gland, which provides temporal control over physiology and behavior (Hatori and Panda 2010). This includes pigment aggregation in melanophores, which express a high affinity melatonin receptor (Ebisawa et al. 1994; Korf et al. 1998). Last, intrinsically photosensitive melanopsin-containing photoreceptors in the retina and pineal complex regulate the background adaptation response by regulating other hormones including α-melanocyte-stimulating hormone (α-MSH) and melanin-concentrating hormone (MCH). A pathway called the retino-suprachiasmatic is involved in background adaptation, with differing amounts of light reaching photoreceptors in the retina affecting the synthesis and secretion of hormones by the pineal gland. For example, tadpoles placed on a black background have less light reaching melanopsin-containing photoreceptors in the retina, which increases secretion of α-MSH and leads to a darkening of the skin (Eberle 1989). Tadpoles placed on a white background exhibit more pigment aggregation due to an inhibition of α-MSH secretion (Tuinhof et al. 1994). Potentially relevant into our ongoing investigation into the mechanisms by which maltol induces melanophore changes, studies have shown that thyrotropin-releasing hormone (TRH) can stimulate the release of MSH in white background-adapted animals, with no effect on animals adapted to a black background (Lidy Verburg-Van Kemenade et al. 1987). In *Xenopus*, melatonin release induces aggregation of melanophores while α-MSH release induces dispersion of melanosomes in melanophores making the skin appear darker (Karlsson et al. 2000).

There has been relatively little research on the effects of chemical toxicants on pigment aggregation in melanophores, especially *in vivo*. However, a study investigating chemical contaminants in water was conducted in cultured *Xenopus* melanophores, where they utilized these cells as a way to screen for potential toxins in water (Iuga et al. 2009). They found that this cell line exhibited a rapid melanosome dispersion response when exposed to 6 out of the 12 chemicals tested, suggesting that a melanophores might be implemented for use as a way of quickly assessing chemical toxicity in drinking water. The Environmental Protection Agency’s Toxicology in the 21^st^ Century (Tox21) program has identified dozens of compounds as both putative thyroid hormone (TH) and retinoid X receptor (RXR) agonists, and maltol, a sugar alcohol that is commonly used as a fragrance and flavoring additive in consumer products and foods, was identified as one of these agonists. In the course of experiments to assess the effects of maltol on TH-dependent mechanisms of neurodevelopment in *Xenopus* tadpoles, we observed that maltol decreased apparent pigmentation, with animals appearing increasingly lighter with increasing concentrations of maltol. The current paper tests potential environmental and hormonal mechanisms by which maltol may exert changes in melanophore pigment aggregation in *Xenopus* tadpoles.

## METHODS

### Animals

We used wild-type *Xenopus laevis* tadpoles bred on site. Hatching was completed in either 20- or 40-gallon tanks kept at 18°C under 12:12 L:D photoperiod. We used stage 48 tadpoles (10-12 days post-fertilization), which were selected based on morphological characteristics in accordance with Nieuwkoop and Faber’s *Xenopus* table of development (1956, 1994). Once selected, tadpoles were moved to glass bowls (200 ml each) in Steinburg’s rearing solution. We fed tadpoles once per day starting when yolk was depleted (∼7 days post-fertilization). We kept tadpoles in a 22°C incubator with 12:12 L:D cycle. Light was 6 inches from the tops of the bowls overhead (Utilitech 65W Equivalent Soft White BR30 LED Flood Light Bulbs).

All animal procedures were performed in accordance with Virginia Tech’s Institutional Animal Care and Use Committee’s rules and regulations.

### Euthanasia and Fixation

Tadpoles were euthanized by 0.2% of tricaine methanesulfonate (MS-222; Sigma, St. Louis, MO, USA) overdose then fixed in 4% phosphate-buffered paraformaldehyde overnight at 4°C prior to imaging.

### Experiment 1: Thyroid hormone and maltol treatments

To systemically treat tadpoles with the thyroid hormone thyroxine (T_4_), we first dissolved 100 µg of crystalline T_4_ (Sigma) in 6.66 ml of 50 mM NaOH (1.93 mM) and further diluted it to 1.93 µM in Steinburg’s solution to make a stock solution, which was stored at −20°C. Aliquots of 1.93 µM T_4_ stock were thawed and 2 mL were added to 198 mL of Steinberg’s solution for a final concentration of 15 µg/L (19.3 nM). To systemically treat tadpoles with maltol, we dissolved 0.1261 g of crystalline maltol (Sigma) in 50 mL warm water to make a stock solution (20mM). For experiment 1, we treated tadpoles systemically with 100 µM, 300 µM, 600 µM, and 1 mM concentrations of maltol, by diluting 1 mL, 3 mL, 6 mL, and 10 mL of the 20 mM maltol stock in 199 mL, 197 mL, 194 mL, and 190 mL of Steinberg’s solution, respectively. Once bath solutions were constituted, approximately 15 tadpoles were placed in bowls for the duration of the respective experiments. Tadpoles were treated for four days then euthanized and fixed.

### Experiment 2: White background vs. Black background

Tadpoles were treated with maltol the same way as in Experiment 1. For tadpoles placed on a white background, white pieces of paper were used to cover the shelf under the bowls for the duration of the experiment. For tadpoles placed on a black background, we placed a specially cut solid black plastic liner to cover the shelf under the bowls for the duration of the experiments. Tadpoles were treated for four days then euthanized and fixed.

### Experiment 3: Melatonin and maltol time course

To treat tadpoles with melatonin, we dissolved 50 mg of melatonin (ChemCruz, Santa Cruz Biotechnology Inc, Dallas, TX, USA) in 50 mL of 100% molecular grade EtOH to create a stock concentration (1 mg/mL). We then diluted 5 µL of stock solution in 200 mL of Steinberg’s solution for a final concentration of 5 ng/mL melatonin. Tadpoles were also treated with 1mM maltol, prepared with methods described above. 5 µL EtOH was added to maltol and control bowls. Tadpoles were treated for four days then euthanized and fixed.

### Imaging

Fixed animals from each group were placed in a custom imaging chamber made from Sylgard, covered with a coverslip, and imaged using transmitted light on a Leica SP8 confocal microscope. Three z-stacks were taken per animal: one of the melanophores in the dura mater over the brain, and one on each side of the brain of the interocular space between the brain and each eye for analysis of dermal melanophores. Images were taken using a 5X (5.25) dry objective with a zoom of 1.5, a z-step size of 15 microns and 5 steps for the images of the brain and a z-step size of 16 microns and 20 steps for the images of the interocular space.

### Quantitative real-time PCR

In order to quantify changes in expression of mchR and POMC, two genes known to be involved in signaling pathways that affect melanophore behavior, we placed stage 48 tadpoles in a 2×2 experimental paradigm bath. In one dimension, tadpoles were treated with either 1 mM maltol or CNTL, in the other dimension, tadpoles were placed over either a white or black background. After four days, we euthanized the tadpoles by overdose of MS-222 (0.2%) and quickly dissected out the brains and placed them into Trizol (Life Technologies), with three brains per tube, and froze them in −80°C. We extracted RNA in accordance with the protocol provided by the manufacturer for Trizol, and performed RNA clean up when necessary. Following extraction, RNA concentrations were measured on a NanoDrop (Thermo Scientific). RNA samples were then reverse transcribed using the Bio-Rad iScript kit using 500 ng of RNA per reaction. We then performed quantitative PCR (qPCR) using 2 ng of cDNA for each reaction using the iTaq Universal SYBR Green Supermix kit (Bio-Rad) on a Bio-Rad CFX384 thermocycler. Primers used for quantification of mchR expression can be found in Table 1.

**TABLE 1.**
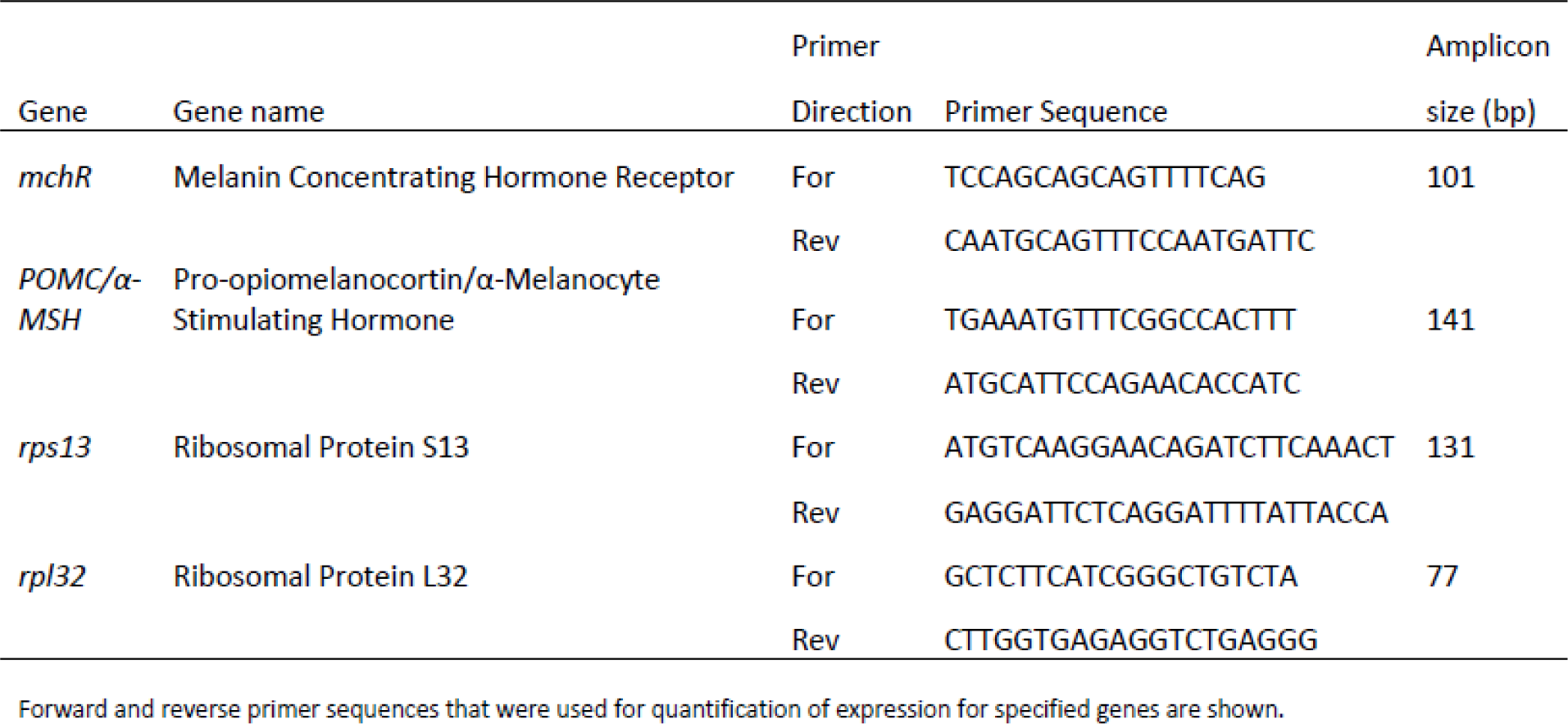
Primers used for quantification of gene expression

All reactions were done in triplicate and outliers with deviations more than 1.5 times the standard deviation from the mean within a set of triplicates were removed from analysis.

## RESULTS

### Maltol induces pigment aggregation in dura mater melanophores in a dose-dependent manner

To evaluate whether maltol induces pigment aggregation in melanophores in the dura mater, we exposed stage 48 tadpoles to CNTL (Steinberg’s rearing solution), T_4_ 15 µg/L, 100 µM maltol, 300 µM maltol, 600 µM maltol, and 1 mM maltol bath for four days. We then imaged the brains of the animals using transmitted light on a confocal microscope and quantified the proportion of the brain that was covered with melanophore pigment. Melanophores in CNTL and T_4_ treated animals were similarly dispersed (Fig. 1A and B). While there appears to be a slight difference in melanophore appearance, with T_4_ appearing to make melanophores appear rounder, these differences were not quantified because melanophore density is so high that most melanophores with dispersed pigment overlap, impeding quantification of individual melanophores. Maltol, on the other hand, induced pigment aggregation of dura mater melanophores in a dose-dependent manner, with pigment aggregated in the center of the melanophore, appearing to be very round (Fig. 1C). Tadpoles treated with 300 µM, 600 µM, and 1 mM concentrations of maltol had a significantly lower proportion of their brains covered with pigment relative to CNTL (Fig. 1D; p<0.0001; one way-ANOVA/Dunnett’s multiple comparisons test).

**Figure 1.**
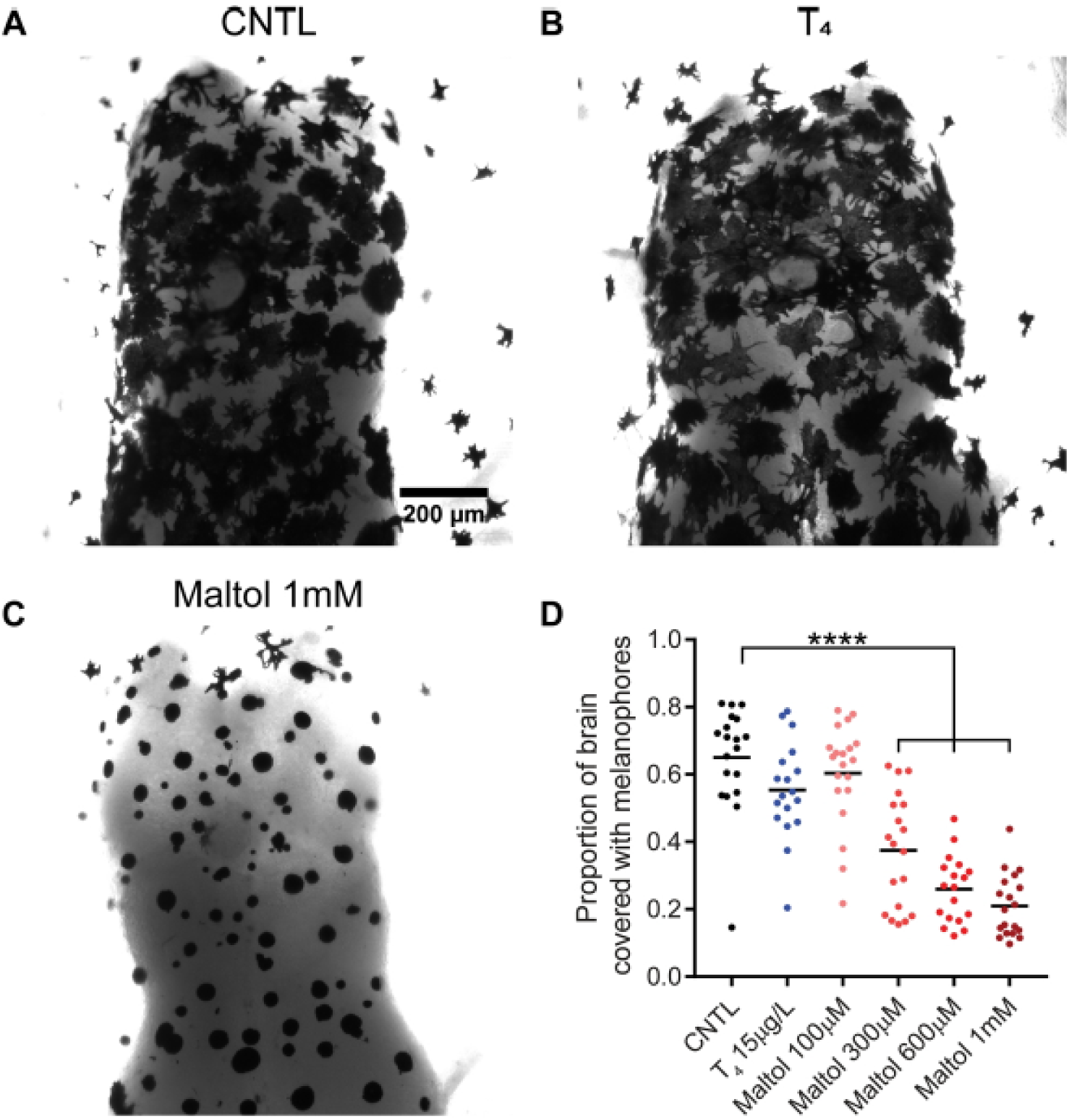
Maltol aggregated pigment in dura mater melanophores in a dose dependent manner. A-C) Representative images illustrating change in pigment aggregation in dura mater melanophores in the telencephalon and optic tectum in A) a control (CNTL) tadpole, B) tadpole treated with T4 for 4 days, and C) tadpole treated with ImM maltol for 4 days. D) Maltol significantly decreased the proportion of the brain surface covered with dura mater melanophore pigment relative to CNTL in a dose dependent manner. Treatment with T4 15pg/L had no effect relative to CNTL. ****p<0.0001

### Maltol induces pigment aggregation in dermal melanophores in a dose-dependent manner

In order to assess the effects of maltol on melanophore pigment aggregation on a cell-by-cell basis, we quantified pigment distribution in dermal melanophores in the area between the brain and each eye. Unlike in the dura mater, the density of dermal melanophores is low enough to allow for measurement of individual melanophores. Nevertheless, the patterns observed in dermal melanophores are similar to those seen in the dura mater. To evaluate degree of pigment aggregation, we quantified melanophore circularity, roundness, and cross-sectional area. Increased circularity and roundness are indicative of aggregation, and increased area of dermal melanophores is consistent with dispersion. As in the dura mater, dermal melanophores in both CNTL and T_4_ treated animals were similarly dispersed and that melanophore shape in these treatment groups were also similar (Fig. 2A and B). 1mM maltol induced substantial aggregation in dermal melanophores relative to CNTL (Fig. 2C). Maltol induced a dose-dependent increase in circularity (Fig. 2D; p=0.0056; p<0.0001; one-way ANOVA/Dunnett’s multiple comparisons test) and roundness (Fig. 2E; p<0.0001; one-way ANOVA/Dunnett’s multiple comparisons test) of dermal melanophores relative to CNTL. For melanophore cross-sectional area, T_4_ increased area relative to CNTL (Fig. 2F; p=0.0493; one-way ANOVA/Dunnett’s multiple comparisons test). Maltol did not significantly affect melanophore area, but a downward non-statistical trend with dose was evident. These results show that 4 days of maltol treatment induces pigment aggregation in a dose-dependent manner, whereas the effect of T_4_ is slight but appears to be opposite to that of maltol.

**Figure 2.**
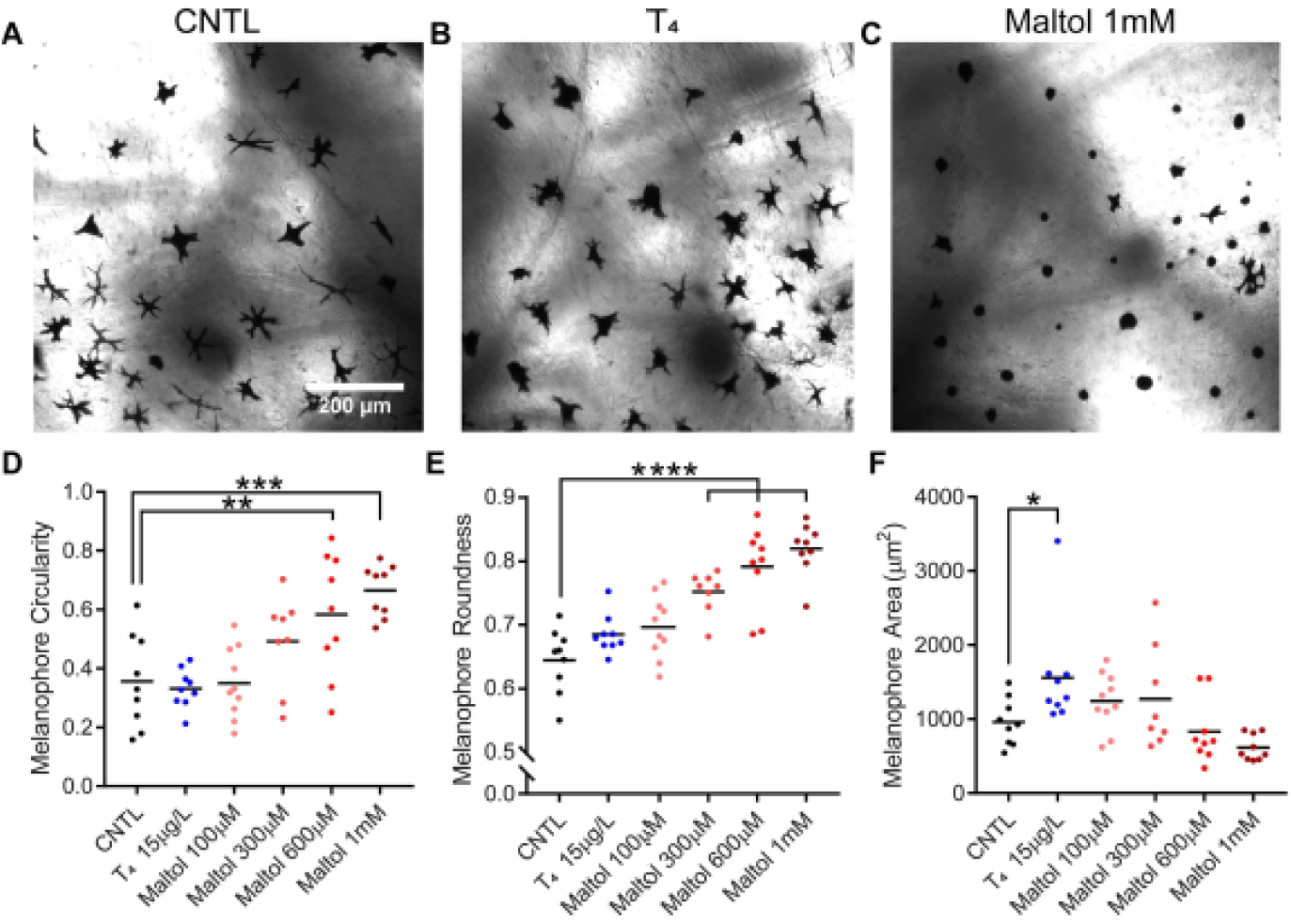
Maltol aggregated pigment in dermal melanophores in a dose dependent manner. A-C) Representative images of dermal melanophores on the dorsal surface of the head between the eye and the brain in A) a CNTL tadpole, B) a tadpole treated with 15 pg/L of T4 for 4 days, and C) a tadpole treated with 1 mM maltol for 4 days. Maltol increased pigment D) circularity and E) roundness in dermal melanophores in a dose dependent manner. F) Maltol did not affect the area of dermal melanophores; T4 significantly increased melanophore area relative to control. *p<0.05, **p<0.005, ***p<0.0005, ****p<0.0001

### Maltol induces pigment aggregation in dura mater melanophores irrespective of background

To ascertain whether rearing tadpoles on black or white backgrounds affects the ability of maltol to induce changes in dura melanophore aggregation, we exposed NF stage 48 tadpoles to CNTL (Steinberg’s rearing solution), 100 µM maltol, 300 µM maltol, 600 µM maltol, and 1 mM maltol bath for four days. White background induces pigment aggregation, whereas black background induces pigment dispersion; background adaptation is mediated by photoreceptors in the retina (Figure 3A). As expected, pigment aggregation in CNTL groups was consistent with the predicted background adaptation response (Fig. 3B). Maltol induced changes in melanophore pigment aggregation and conformation irrespective of background, with the highest observed degree of aggregation in animals reared on a white background and in 1 mM maltol bath. Maltol induced a dose-dependent decrease in proportion of the brain covered with pigment in both white and black background groups (Fig. 3C). Specifically, 1 mM maltol treatment groups had significantly decreased pigmented brain surface area proportions relative to controls regardless of background (black: p=0.0404, p<0.0001 white: p=0.0038, p<0.0001; 2-way ANOVA/Dunnett’s multiple comparisons test). We also normalized the data to CNTL groups for each background condition in order to observe the effect of maltol irrespective of background (Fig. 3D). Maltol appeared to induce a similar degree of change with dose, with the only statistically significant difference seen between the black and white background 1 mM maltol groups (p=0.0094; 2-way ANOVA/Sidak’s multiple comparisons test). These data show that maltol induced changes in dura melanophores with increasing concentrations regardless of background condition.

**Figure 3.**
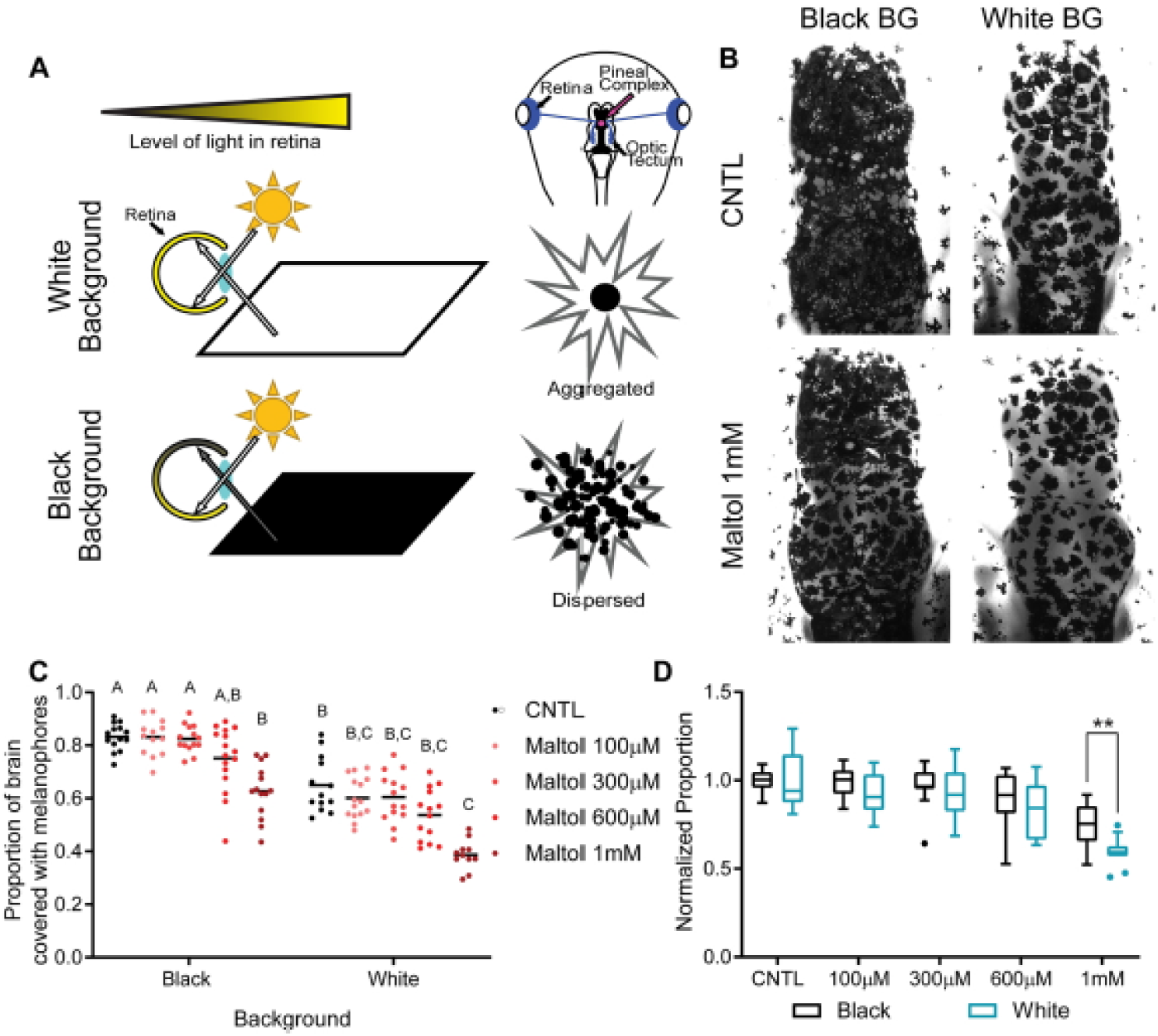
Maltol induces similar patterns of pigment aggregation in dura mater melanophores in tadpoles over white or black backgrounds. A) Schematic depicting background adaptation mechanism in Xenopus laevis tadpoles, where pigment aggregation is regulated by the reflection of light onto the dorsal retina. B) Representative images of pigment aggregation in dura mater melanophores in CNTL tadpoles or tadpoles exposed to 1 mM maltol for 4 days on either a white or black background (BG). C) Maltol decreased the proportion of the brain covered with pigment in a dose dependent manner in both white and black backgrounds. D) Data in C normalized to CNTL to illustrate that the dose dependent effect of maltol was largely independent upon background except at the highest concentration. **p<0.005, ****p<0.0001

### Maltol induces changes in dermal melanophore shape irrespective of background

To evaluate whether rearing tadpoles on black or white backgrounds influences the ability of maltol to induce pigment aggregation in individual melanophores, we again examined dermal melanophores in the area between the brain and each eye in the same animals as in Fig 3. We again saw differences in interocular dermal melanophore pigment distribution that were consistent with predicted background adaptation responses (Fig. 4A). Specifically, dermal melanophores of animals from treatment groups reared on a black background showed a higher degree of dispersion relative to their white background counterparts. Maltol increased circularity in a dose-dependent manner across both backgrounds. Normalized circularity to control groups revealed significant differences between the black and white background groups for both 600 µM and 1 mM maltol treatment groups (Fig. 4C; p=0.002; p<0.0001; 2-way ANOVA/Sidak’s multiple comparisons test). Maltol also appeared to increase roundness of dermal melanophores in a dose-dependent manner, but the only statistically significant difference in this measure was between the white background control and 1 mM maltol groups (Fig. 4D; p=0.0073; 2-way ANOVA/Dunnett’s multiple comparisons test). Normalizing the values to the controls for each group showed the general upward trend with increasing concentrations, and there were no significant differences between backgrounds (Fig. 4E). Maltol decreased pigment distribution in dermal melanophores in a dose-dependent manner in both backgrounds (Fig. 4F). For the black background groups, statistically significant differences began at the 300 µM maltol concentration (p=0.0126; 2-way ANOVA/Dunnett’s multiple comparisons test) and were also seen for both the 600 µM and 1 mM concentrations (p<0.0001; 2-way ANOVA/Dunnett’s multiple comparisons test). For the white background groups, significant differences were seen in the 300 µM and the 1 mM maltol concentrations (Fig. 4F; p=0.0268; p=0.0172; 2-way ANOVA/Dunnett’s multiple comparisons test). Normalized area relative to CNTL illustrated the general downward trend with increasing concentrations, and there were no significant differences between backgrounds (Fig. 4G). Collectively, the dose-dependent increases in melanophore circularity and roundness as well as the dose-dependent decreases in melanophore area show that maltol induces pigment aggregation regardless of rearing background.

**Figure 4.**
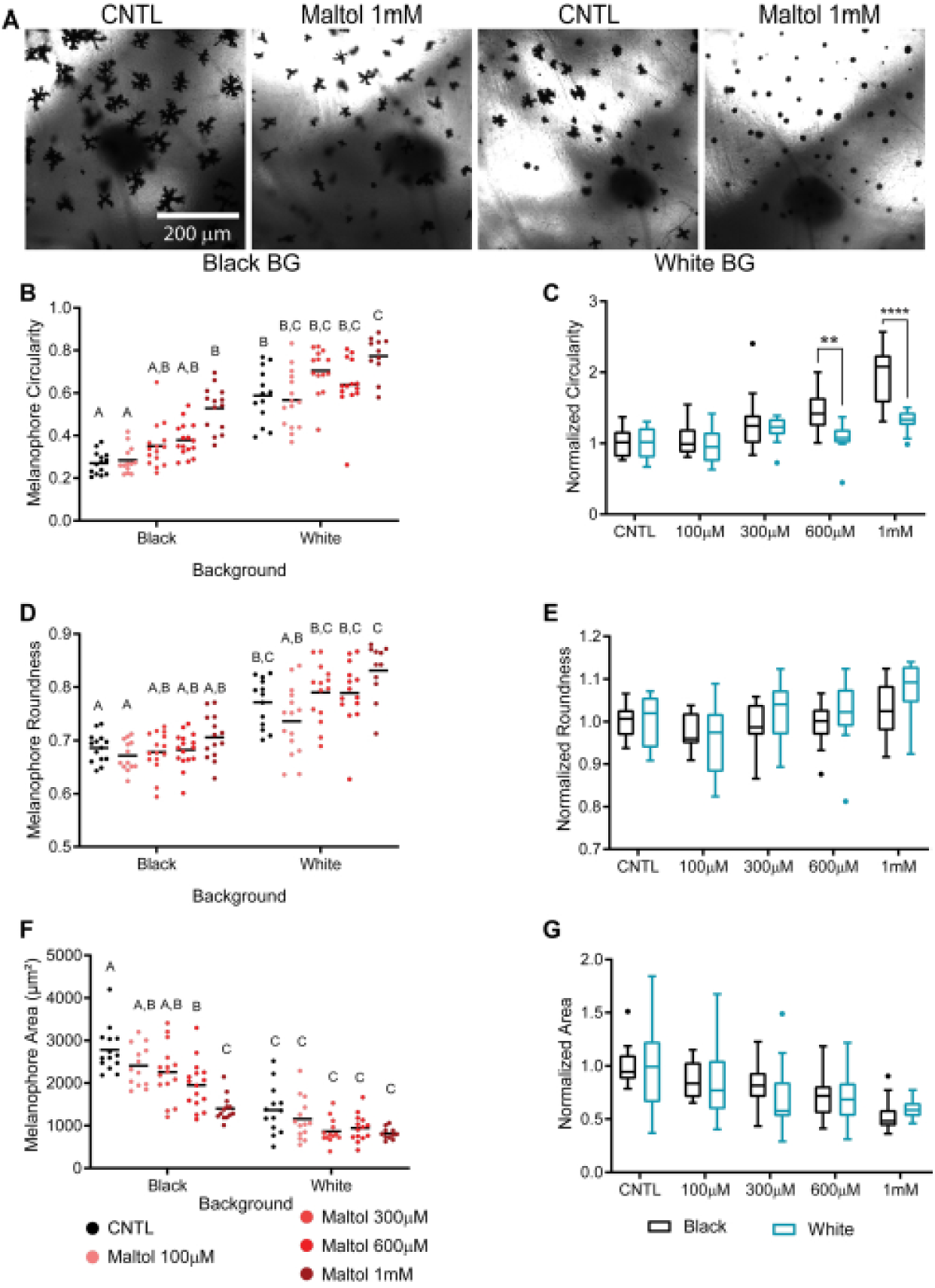
Maltol induces similar patterns of pigment aggregation in dermal melanophores in tadpoles over white or black backgrounds. A) Representative images of dermal melanophores in both CNTL tadpoles and tadpoles treated with 1 mM maltol reared on either a black or white background. Maltol increased B) circularity and D) roundness, and decreased F) average area of dermal melanophores in a dose dependent manner. C, E, G) Data in B, D, F, respectively, normalized to CNTL to illustrate that the dose dependent effect of maltol on pigment aggregation was largely independent upon background except at the highest concentrations. *p<0.05, **p<0.01, ***p<0.0005, ****p<0.0001

### Maltol does not induce changes in MCHR and α*-MSH expression*

To assess whether maltol exposure may affect pigment distribution in melanophores by changing expression of hormones known to affect pigment distribution and conformation in *Xenopus*, we exposed NF stage 48 tadpoles on black or white backgrounds to either CNTL or 1 mM maltol bath for four days. We then dissected out the brains and used qPCR to evaluate expression of both melanin concentrating hormone receptor (MCHR) and α-melanocyte stimulating hormone (α-MSH). These hormones both affect melanophore behavior, with MCH typically being involved in inducing aggregation and α-MSH secretion involved in dispersion of melanophores. Quantification of gene expression revealed no discernible interaction, background, or treatment effect on POMC expression (Descriptive statistics shown in Table 2; F_(1,20)_= 1.532 p=0.2302, F_(1,20)_= 2.109 p=0.1620, F_(1,20)_= 0.1285 p=0.7238, respectively; 2-way ANOVA). Quantification of gene expression also revealed no discernible interaction, background, or treatment effect on mchR expression (Descriptive statistics shown in Table 2; F_(1,19)_= 0.09621 p=0.7598, F_(1,19)_= 0.2848 p=0.5998, F_(1,19)_= 0.5198 p=0.4797, respectively; 2-way ANOVA). Further, Sidak’s multiple comparisons test for both POMC and MCHR revealed no statistically significant differences either between treatment groups or rearing background. These data show that maltol–mediated changes in pigment aggregation is likely not via changes in expression of either POMC or MCHR.

**TABLE 2.** Quantification of Expression of Melanophore Regulating Hormones^1^

|  | Mean Estimated Expression |  |  |  |  |  |
| --- | --- | --- | --- | --- | --- | --- |
| Treatment | mchR | | POMC/α-MSH | | | |
| Black Background |  |  |  |  |  |  |
| CNTL | 1.0 ± 0.12 | | 1.08 ± 0.17 | | | |
| Maltol 1 mM | 0.88 ± 0.17 | | 0.74 ± 0.18 | | | |
| White Background |  |  |  |  |  |  |
| CNTL | 1.1 ± 0.15 | | 1.04 ± 0.12 | | | |
| Maltol 1 mM | 1.0 ± 0.16 | | 1.24 ± 0.24 | | | |
|  | Descriptive Statistics |  |  |  |  |  |
| | mchR | | | POMC/α-MSH | | |
|  | df | f | p | df | f | p |
| Background Effect | 19 | 0.2848 | 0.5998 | 20 | 2.109 | 0.162 |
| Treatment Effect | 19 | 0.5198 | 0.4797 | 20 | 0.1285 | 0.7238 |
| Interaction Effect | 19 | 0.09621 | 0.7598 | 20 | 1.532 | 0.2302 |

### Maltol induces dynamic changes in both dura and dermal melanophores over time

To establish the timing of maltol-induced melanophore changes, we conducted a time course experiment where we exposed NF stage 47-48 tadpoles to either CNTL (Steinberg’s rearing solution), melatonin 5 ng/mL, or 1 mM maltol bath for 48 hours. We sacrificed and fixed 6 or more animals at 8 time points: 0 minutes, 15 minutes, 30 minutes, 1 hour, 2 hours, 6 hours, 24 hours, and 48 hours. Because we know that melanophores are dynamic in response to light, we chose the first time point at lights on in the morning. We used melatonin 5 ng/mL as a positive control, as it has been well established by prior studies to regulate melanophore behavior in *Xenopus*, inducing pigment aggregation during night time (Binkley et al. 1988). Melatonin significantly decreased the proportion of the brain covered by melanophores at the 15-minute, 30-minute, and 6-hour time points when compared with controls (Fig. 5A&B; p<0.0001; p=0.0024; 2-way ANOVA/Tukey’s multiple comparisons test). Maltol did not significantly decrease proportion of the brain covered with pigment relative to CNTL during the 48 hr of treatment. Melatonin significantly decreased melanophore area at 30-minute, 1-hour, and 24-hour time points relative to controls (Fig. 5C; p=0.0169; p=0.0446; p=0.0419; 2-way ANOVA/Tukey’s multiple comparisons test). Conversely, maltol appears to initially induce dispersion of pigment in melanophores at earlier time points up to and including 2 hours of exposure, with a significant difference in melanophore area between 1 mM maltol and CNTL (Fig. 5A&C; p=0.0411; 2-way ANOVA/Tukey’s multiple comparisons test). Following the 2-hour time point, average melanophore area in the maltol treatment group trended downward. Melatonin increased melanophore circularity in the melatonin treatment group at 15 minutes, 30 minutes, 1 hour, 6 hours, and 48 hours relative to controls (Fig. 5D; p<0.0001; p<0.0001; p=0.0157; p=0.0266; p=0.0345; 2-way ANOVA/Tukey’s multiple comparisons test). Maltol significantly increased circularity at the 48 hr time point (Fig. 5D; p<0.0001; 2-way ANOVA/Tukey’s multiple comparisons test). Roundness was similar to circularity, with significant differences between melatonin and control groups at 15 and 30 minutes and between maltol and control groups at the 48-hour time point (Fig. 5E; p<0.0001; 2-way ANOVA/Tukey’s multiple comparisons test). Together, these data demonstrate that maltol induces dynamic changes in both dura and dermal melanophores, but that time course of pigment aggregation is dissimilar to treatment with 5 ng/mL melatonin.

**Figure 5.**
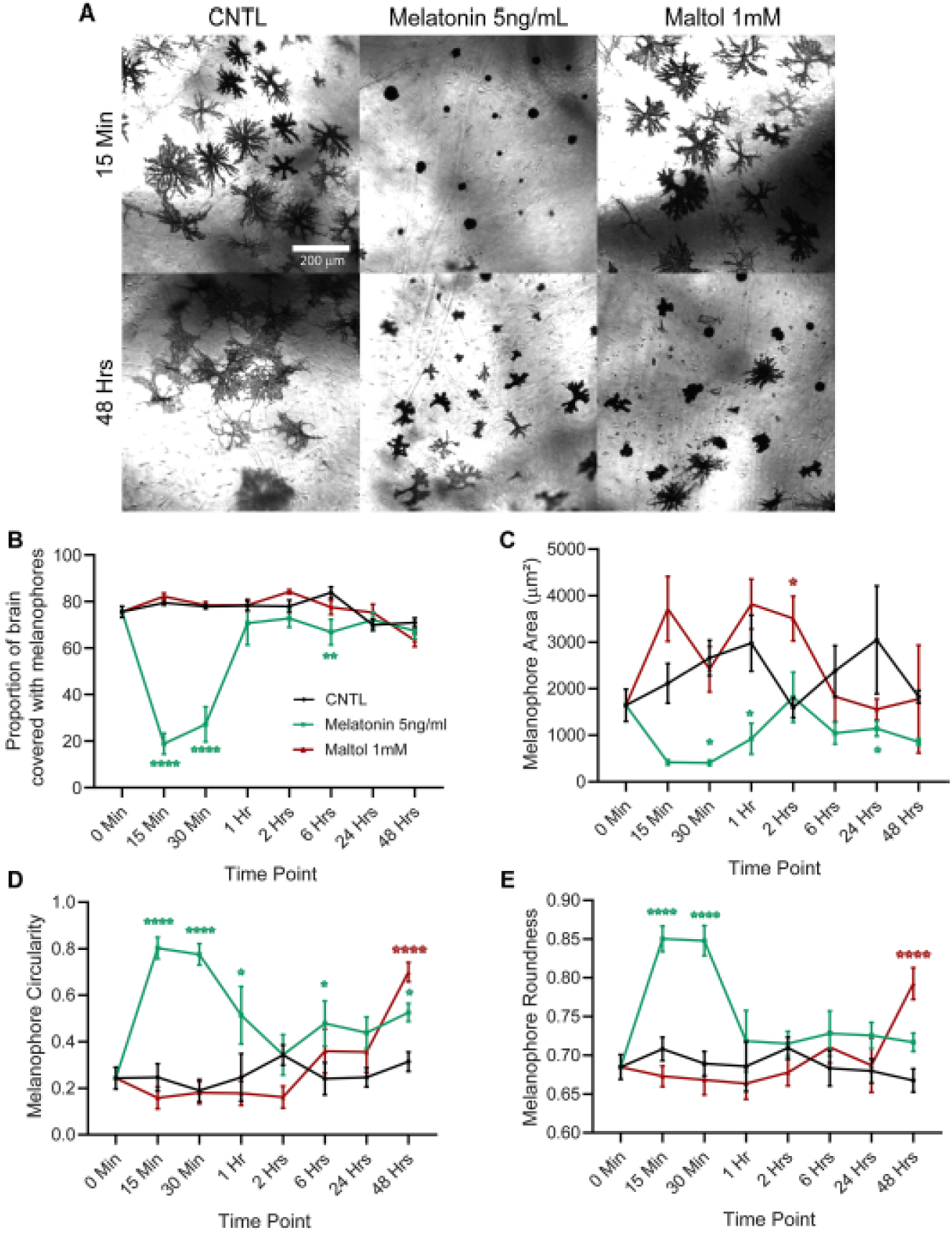
Melatonin induced pigment aggregation within 15 min, whereas maltol induced pigment aggregation at 48 hrs post treatment onset. A) Representative images of dermal melanophores in CNTL, melatonin (5ng/mL), and maltol (1 mM) treated animals at 15 minutes and 48 hours. B) Melatonin induced temporary pigment aggregation in dura mater melanophores within 15 min; pigment was dispersed by 1 hr. Maltol did not induce pigment aggregation in dura mater melanophores by 48 hrs. C) Melatonin significantly decreased average dermal melanophore area within 30 min relative to CNTL. Maltol significantly increased dermal melanophore area at 2 hrs, and only marginally increased melanophore are over the few hours. Melatonin significantly increased D) circularity and E) roundness of dermal melanophores within 15 min, whereas maltol increased circularity and roundness at 48 hours. *p<0.05, **p<0.01, ****p<0.000 Note: Green asterisks denote significance between CNTL and melatonin treated animals; red asterisks denote significance between CNTL and maltol treated animals.

## DISCUSSION/CONCLUSIONS

Our results show that maltol induced pigment aggregation in both dura and dermal melanophores in the *Xenopus laevis* tadpole. Furthermore, maltol induced melanophore aggregation in a way that is independent of the background adaptation mechanism present in these animals. Finally, maltol induced pigment aggregation after 48hr treatment, which was much slower than the effects of melatonin. Collectively, these data show that systemic exposure to maltol leads to changes in pigment distribution in the form of aggregation of melanophores in both the dura mater over the brain and in the skin of *Xenopus laevis* tadpoles but that maltol’s effects are likely not mediated by the background adaptation mechanism nor by changes in melatonin signaling.

We found that maltol affects pigment aggregation similarly under white and black background, suggesting that the maltol does not act on melanophores via the background adaptation mechanism. Superficially, normalized results from Experiment 2 suggest that there may be subtle differences in maltol’s response across backgrounds (Fig 3D and Fig 4C). These differences may be explained by the fact that the nature of the data is bounded, i.e. limited to values between zero and one. Given that data were normalized to the CNTL group and that pigment aggregation circularity under white background is greater than 0.5, it is impossible to have circularity values that could approach the normalized values under black background. Regardless, the normalized measures reveal very similar trends in each measure induced by exposure to increasing concentrations of maltol. Multiple studies recently have characterized the role of melanopsin-containing retinal ganglion cells in dynamic changes in skin pigmentation. For example, it was demonstrated that activation of these RGCs block α-MSH secretion by inhibition of melanotropes mediated by GABA_A_R (Bertolesi et al. 2016). They also demonstrated that melanopsin-containing RGC activation leads to changes in skin pigmentation resulting in the animal appearing lighter, and that this is mediated by a secondary response that negatively regulates α-MSH secretion (Bertolesi et al. 2015). Future experiments could further explore this issue by exposing eyeless tadpoles to maltol to ascertain whether the same maltol-induced pigment aggregation occurs independent of input to non-image forming photoreceptors in the eye. If eyeless tadpoles exhibit the same melanophore changes in response to maltol exposure, it would suggest that maltol is inducing these changes at the level of the melanophores themselves or through another unknown mechanism.

Changes in melanophore pigment distribution in many animals are mediated primarily by two hormones secreted by the brain’s pituitary gland, α-melanocyte stimulating hormone (α-MSH) and melanin concentrating hormone (MCH). Release of α-MSH typically induces melanophore dispersion, whereas MCH is typically responsible for melanophore aggregation. Tadpoles placed on a black background have an increase in circulating α-MSH resulting in darkening of skin color (Eberle 1989). Conversely, placing animals on a white background inhibits α-MSH secretion and skin color appears lighter because of pigment aggregation. This mechanism is regulated by a retino-suprachiasmatic pathway that controls melanotrope cells in the pituitary (Tuinhof et al. 1994), which are responsible for synthesizing and secreting α-MSH by cleaving it from its precursor pro-opiomelanocortin (POMC). We initially hypothesized that maltol was either downregulating α-MSH or upregulating MCHR as a mechanism to induce changes in melanophores. We found that our qPCR data evaluating expression of both MCHR and POMC did not show any significant differences induced by maltol between groups on black and white backgrounds. While our data suggest that the mechanism by which maltol induces pigment aggregation seems to be independent of these two hormone systems, these data are not conclusive, however. A limitation of this particular part of this study is that qPCR may not fully capture dynamic changes in circulating levels of α-MSH since it is cleaved from precursor peptide POMC. Our primers targeted POMC mRNA, and posttranslational regulation of POMC transcript to generate α-MSH would not be quantified in our approach. A previous study used radioimmunoassy to show in *Xenopus* that the pars intermedia of the pituitary gland synthesizes and stores α-MSH for later release when on a white background and secretes it when animals are moved to a black background (Jenks et al. 1977). Other studies on this phenomenon utilized immunocytochemistry to quantify changes in α-MSH (Kramer et al. 2003; Roubos et al. 2010). Additionally, we would not expect to see much of a change in MCH as the background adaptation response in amphibians is primarily dependent on regulation of α-MSH by melanopsin-expressing photoreceptors, whereas MCH is the main hormonal regulator of pigment distribution in teleost melanophores (Bertolesi and McFarlane 2018). Although more research will conclusively determine whether maltol affects pigment distribution via alterations in the neural and hormonal mechanisms that control background adaptation, our data strongly suggest that maltol does not act via the background adaptation mechanism.

Melatonin-induced pigment aggregation is a well-established phenomenon and has been documented in multiple animals, including both amphibians and fish (Filadelfi et al. 1994). In fact, melatonin gets its name from its initial discovery in feeding bovine pineal gland to tadpoles, which “toned” the melanin and induced pigment aggregation. The behavior of melanophores in the melatonin-treated animals over time reaffirm results from previous studies investigating melatonin-induced melanophore changes. For example, melatonin and melatonin analogues induce robust pigment aggregation in melanophores quickly, on the order of minutes (Sugden 1992). Interestingly, cultured *Xenopus* melanophores exhibit melatonin-induced desensitization, where the induction of aggregation no longer occurs in the same way with prolonged exposure to melatonin (Rollag and Lynch 1993). This is consistent with what we observed in the time course experiment, where animals were systemically treated with melatonin for the duration of the experiment. Specifically, melatonin-treated animals showed quick and drastic pigment aggregation until the 30-minute time point but began to disperse after that and remained similar to that of control- and maltol-treated animals for the remainder of the experiment. Maltol, on the other hand, did not induce sudden pigment aggregation. Instead, pigment aggregation was not noticeable until 48 hr of treatment, long after the effects of melatonin have worn off. Furthermore, results from Fig. 1 show that maltol aggregated pigment at the 4 day time point, and therefore the effects of maltol emerge more slowly and are much more long lasting than melatonin, lasting at least two days once aggregated. These results suggest that maltol likely does not act via a melatonin-based mechanism, such as binding to melatonin receptor or stimulation of melatonin from the pineal gland. Nevertheless, pinealectomy would help rule out if maltol does induce pigment aggregation via activation of the pineal gland.

These results provide insight to toxicity of a particularly sensitive cell type that is influenced via a wide range of stimuli and offer the opportunity to better understand how some environmental chemicals may impact overall physiology through specific sub-cellular mechanisms. It is important to note, however, that there are limitations in relating these findings to human health. First, the route of administration in this study differs from the way that humans would be exposed to maltol. These animals were treated via bath exposure, whereas the most common route of exposure for humans is through ingestion of maltol-containing foods. That said, there are topical application of maltol-containing fragrances and products which may allow for maltol to be absorbed through the skin, which is somewhat similar to our route of administration. Furthermore, maltol is now an ingredient in e-cigarettes, and therefore absorbance via inhalation into the bloodstream shares important similarities to bath application of maltol, which may enter tadpoles via the gills. Another important caveat is that the concentrations used in these studies are likely to be higher than what humans are exposed to at any given time, but the degree to which people are exposed to maltol in their environments is not well characterized. Exposure predictions from the EPA Chemistry Dashboard predict 95 percentile exposure levels that are no more than 1.7 ug/kg bodyweight per day (Williams et al. 2017). The logP for maltol (−0.276) suggests that it does not bioaccumulate and likely expelled relatively quickly. Thus, although our data show that maltol is capable of affecting this sensitive cell population in striking ways, much more work needs to be done before drawing conclusions about the potential effects of maltol on human health.

Our results strongly suggest that maltol does not affect melanophore pigment aggregation via the background adaptation or the melatonin mechanisms, and therefore the mechanisms by which maltol affects these changes have yet to be elucidated. Future experiments could evaluate several other potential mechanisms that maltol may act on to induce the changes described here. First, the melanophores themselves are photosensitive as they express melanopsin to mediate changes in melanophore behavior to fulfill needs like thermoregulation and UV protection (Bertolesi and McFarlane 2018). Maltol may affect melanophore melanopsin physiology, impairing these cells from dispersing pigment properly when exposed to light. One potential mechanism was described in studies done in cultured *Xenopus* melanophores, where it was found that treatment with all-trans retinal resulted made melanophores photosensitive and affected the degree to which pigment dispersed when exposed to light (Rollag 1996). Given that maltol has been identified as a putative retinal X receptor agonist in the Tox21 data set, maltol may affect pigment aggregation via a retinoic acid-signaling based mechanism. This is particularly interesting because retinoids are commonly used in skin creams and other cosmetic products to lighten the skin, which would be consistent with the effects of maltol reported here. If maltol affects melanophore pigment aggregation via retinoic acid signaling, then *Xenopus* melanophores may be well suited as a cellular model for initial screening of potential disruptors retinoid-mediated changes in pigmentation.

## ACKNOWLEDGEMENTS

Special thanks to our undergraduate students Robert Bass and Hannah Sturgeon for help in caring for the animals.

## DATA ACCESSABILITY STATEMENT

The data can be made available to readers by contacting the authors of this study.

